# A Wake-Up Call: Determination of Antibiotics Residue Level in Raw Meat in Abattoir and Selected Slaughterhouses in Five Local Government in Kano State, Nigeria

**DOI:** 10.1101/2022.01.04.474991

**Authors:** Habeeb Modupe Lateefat, Opasola Afolabi Olaniyi, Garba Misbahu, Olalekan Morufu Raimi

## Abstract

The frequent use of antibiotics may result in drug residues that can be discovered at varying quantities in animal products such as milk or meat. The presence of pharmaceutical residues in food above the MRLs has been perceived globally by various persons. Antibiotics residues are present in food, which can endanger human health by causing antibiotic sensitivity, allergic reactions, microflora imbalance, bacterial resistance to antibiotics in microorganisms, and financial loss to the food industry. Farmers around the world utilize them on a sporadic basis for both preventative and curative purposes. This study assessed the antibiotics residues in raw meat sold in 6 slaughter houses in Kano States. The study is a descriptive cross-sectional study involving six (6) major slaughter house in Kano state. Muscle, Kidney and liver samples were collected from each slaughterhouse. The antibiotic residues in the meat samples were analysed using high performance liquid chromatography (HPLC) for tetracycline, ciprofloxacin and oxytetracycline residue results were presented in charts and tables. Out of a total of 18 beef samples analyzed during this study, 15 (83%) of the total samples had detectable levels of tetracycline residues from which 6(33.3%) had tetracycline residues at violative levels above the WHO/FAO maximum residue limits (MRLs), out of those 18 beef samples analyzed during this study, 6(33%) of the total samples had detectable levels of oxytetracycline residues from which 3(17%) had oxytetracycline residues at violative levels above the WHO/FAO maximum residue limits (MRLs) and out of those 18 beef samples analyzed during this study, 12(67%) of the total samples had detectable levels of ciprofloxacin, all levels are below the WHO/FAO maximum residue limits (MRLs). This high level of tetracycline and oxytetracycline residues in greater proportion of meat destined for human consumption at violative levels could be as a result of the indiscriminate use and misuse of veterinary drugs as commonly practiced among livestock producers and marketers without observing withdrawal period prior to slaughter. These results indicate that consumers may be predisposed to health hazards and hinder international meat trade from Nigeria. Regulatory authorities should therefore ensure compliance with good agricultural practices including withdrawal period of drugs used for treatment of food animals.

## 1. Introduction

Rachel Carson stated fifty (50) years ago for the first time that the widespread use of pesticides is hazardous not only to wildlife but also to humans. This is still an ongoing concern, as evidenced by recent studies by (Olalekan *et al*., 2020d; Raimi *et al*., 2020b; Hussain *et al*., 2021a; Hussain *et al*., 2021b; Isah *et al*., 2020a; Isah *et al*., 2020b; Morufu *et al*., 2021f; Morufu, 2021; Olalekan *et al*., 2021), that pesticides are contributing to and causing massive adverse effects, necessitating the restriction of a wide range of chemical products and uses, and the use of many chemicals in commerce with far greater prudence and caution. The authors claimed that humankind is heading beyond a “safe operating area” as the size of these consequences approaches or exceeds certain thresholds that indicate global system tipping points or natural limits for processes with no apparent limits (so-called “hazardous levels”). Numerous researchers have reported that human health is generally influenced by their environment (Ecological Security Organization (EPA), 2015), Raimi and Sabinus, 2017; Raimi *et al*., 2017; Olalekan *et al*., 2018; Suleiman *et al*., 2019; Raimi *et al*., 2019a; Raimi *et al*., 2019b; Raimi *et al*., 2019c; Olalekan *et al*., 2019; Okoyen *et al*., 2020; Adedoyin *et al*., 2020; Olalekan *et al*., 2020a; Olalekan *et al*., 2020b; Olalekan *et al*., 2020c; Raimi *et al*., 2020; Raimi *et al*., 2021a; Raimi *et al*., 2021b; Raimi *et al*., 2021c; Morufu *et al*., 2021a; Morufu *et al*., 2021b; Morufu *et al*., 2021c; Morufu *et al*., 2021d; Morufu *et al*., 2021e) and specifically, the nature and quality of the food (Ames, 1983; Oluwaseun *et al*., 2019; Afolabi and Raimi, 2021). Nature cattle products is broadly concerning public health body throughout the globe since veterinary medications have assumed a significant part in the field of animal farming and agro-industry, and increasing event of drug residues, and resistance have become issues of concern (Rokka *et al*., 2005). Veterinary medications are basically expected to address the difficulties in provision of sufficient measures of food to the increasing global population (Crawford, 1985) as medications enhance pace of weight gain, further enhances feed proficiency, or forestall and treat infections in food producing animals (Crawford, 1985 and American Veterinary Medication Affiliation (AVMA), 2015).

Veterinary Anti-infection agents (VAs) are mainly used by cattle rearers and poultry farmers, that might prompt antibiotic residues from food animals to human, and consequently cause dangerous health hazards to the consumers (Chanda *et al*., 2014). High degree of antibiotics residues consumption from animal products to human might affect immunological reactions and can unfavorably influence digestive microbiota in susceptible people (Ramatla *et al*., 2017). Nonetheless, farmers utilize the VAs for various purposes such as prophylactic, healing, growth promoters and at times both prophylactic and healing purposes throughout the globe (Wadoum *et al*., 2016). In Canadian poultry industry for prophylaxis and growth enhancers are used for various types of antibiotics are used (Diarra and Malouin, 2014). Enormous level of antibiotics are utilized in Bangladesh yearly. A large proportion of this are used unreasonably under conditions of lacking or no skilled supervision and as a rule without earlier testing on documentation of the microorganisms and resilience of its sensitivity of the antibiotics recommended (Sarker *et al*., 2016).

Fluoroquinolones including ciprofloxacin, enrofloxacin, nalidixic corrosive, and so on have been largely used for treatment and prophylaxis (emergency room *et al*., 2013). Ciprofloxacin is observed viable where microorganisms are resistant to aminoglycosides, tetracycline, macrolides and β-lactams (Ruler, 2014). Tetracycline serves various function such as treating infections and as a growth enhancer in animals owing to the fact that it is a broad-spectrum antibiotic. Around 60% of an ingested portion of oxytetracycline is assimilated from the gastrointestinal tract and broadly spread in the body (Doyle, 2006; Mund *et al*., 2017). As of late, many examinations have shown that antibiotics injected to poultry and domesticated animals were amassed in liver, kidney, muscle and bones surpassing the Maximum Residual Limits (MRL) (see table 1 below) (Sarker *et al*., 2016). The often use of antibiotics might bring about drug residues that can be found at various levels in animal products like milk or meat. Presence of medications residues in food beyond the MRLs has been perceived worldwide by different people (Kempe and Verachtert, 2000). Presence of antibiotics residues in food stuff can present risks to human wellbeing e.g., sensitivity to antibiotics, allergy responses, microflora imbalance, bacterial resistance to antibiotics in microorganisms and loss to the food business. Notwithstanding, farmer suse them occasionally for both prophylactic and remedial purposes globally (Wadoum *et al*., 2016). Thus, the aim of the research is to aimed at evaluating of chemical level and antibiotic residue in raw meat and having the following objectives which are: to determine the level of chemical and antibiotic residues in meat samples and to compare the level of chemical and antibiotic residues in the different meat from different abattoirs.

**Table 1:** Maximum residual limits (MRLs) of Tetracyclines and Oxytetracycline in animal derived foods according to the WHO, 1999

| S/N | Antibiotic | Tissue | MRLs |
| --- | --- | --- | --- |
| 1 | Tetracyclines (Oxytetracycline, | Muscle | 200 µg/kg |
| 2 | Tetracycline, chlortetracyclines) | Liver | 600 µg/kg |
| 3 |  | Kidney | 1200 µg/kg |

## 2. Materials and Methods

### Research Design

This is a descriptive cross-sectional design that involved laboratory analysis to determine the level of antibiotic residues in meat samples which was based on laboratory analysis.

### Study Area

Kano is the state capital of Kano State in North West, Nigeria. It is situated in the Sahelian geographic region, south of the Sahara. Kano is the commercial nerve centre of Northern Nigeria and is the second largest city in Nigeria. The Kano metropolis initiallycovered 137 square kilometres (53 square miles), and comprised six local govern ment areas (LGAs) Kano Municipal, Fagge, Dala, Gwale, Tarauni and Nasarawa; However, it now covers two additional LGAs - Ungogo and Kumbotso. The total area of Metropolitan Kano is now 499 square kilometres (193 square miles), with a population of 2,828,861 as of the 2006 Nigerian census; the latest official estimate (for 2016) is 3,931,300. The principal inhabitants of the city are the Hausa people. However, there are many who speak Fulani language. As in most parts of northern Nigeria, the Hausa language is widely spoken in Kano. The city is the capital of the Kano Emirate.

### Field Sample Collection

**Table 2:** Location and Number of Samples Collected from Kano Abattoir and other Slaughter House within the state.

| LOCATION | SAMPLE |
| --- | --- |
| Kano Abattoir Limited | 1 |
| Unguwa Uku Slaughter House | 2 |
| Dawanau Slaughter House | 3 |
| Badarawa Slaughter House | 4 |
| Madubi Slaughter House | 5 |
| Warawa Slaughter House | 6 |

### Sampling Technique

Kidney, liver and muscle samples were randomly selected from six slaughter houses in Kano state. Totaling 18 samples.

### Sample Collection

A total of 18 samples (kidney, muscle and liver) were collected from Kano abattoir limited and other slaughter houses from different local government of Kano state. Each sample was placed into a separate plastic container. The collected samples were stored in Freezer at −20°C and transferred to laboratory in a cooler of ice and until extraction.

### Apparatus

JASCO HPLC system with pump PU1580, a Rheodyne loop injector with injection volume 20uL, JASCO UV detector 1575 operated at 365nm, JASCO BORWIN software version 1.50 with Hercule-200 chromatography interface for results integration and recording, HIQ Sil C_18_ LC column (main column size: 4.6mm internal diameter x 250mm length and guard column size: 4.6mm internal diameter x 75mm length) and mobile phase made of 50mM aqueous oxalic acid solution containing 13% (V/V) each of methanol and acetonitrile were used.

### Reagent

HPLC grade water filtered through 0.2μm with maximum impurities of 0.0003% and minimum transmission of 100% at 200 and 250 nm procured from Qualigens Fine chemicals, Glaxo-Smithkline Pharmaceutical Ltd was used.

### Extraction

2g of raw meat product was homogenized in a blender for 2 min and then 0.1g citric acid was added. To this mixture, 1ml nitric acid (30%), 4ml methanol and 1ml deionised water were added, respectively. The suspension with solid particles was put in a vortex for good mixing, kept in an ultrasonic bath for 15 min and then centrifuged for 10 min at 5300 rPM. After filtering through a 0.45μm nylon filter, 20μl of solution was injected into HPLC for analysis.

### Data Analysis

The data was analyzed using mean, percentage and chart.

## 3. Results

The result of sample 1 above of 1 slaughtered cattle. A total of 3 tissues (liver = 1; kidney = 1; muscle = 1) were screened for the presence of antibiotics residue. Tetracycline and ciprofloxacin were detected. The mean residue levels of tetracycline were 17.57 μg/kg ± 6.20 μg/kg and 82.77 μg/kg ± 12.60 μg/kg for ciprofloxacin respectively. The result shows multi-residues of antibiotics

It can be observed that from the sample 2 above of 1 slaughtered cattle. A total of 3 tissues (liver = 1; kidney = 1; muscle = 1) were screened for the presence of drugs residue. Only tetracycline was detected. The mean residue levels of tetracycline were 575.37 μg/kg ± 73.23 μg/kg.

From the above samples 3 of 1 slaughtered cattle. A total of 3 tissues (liver = 1; kidney = 1; muscle = 1) were screened for the presence of antibiotics residue. Tetracycline, ciprofloxacin and oxytetracycline were detected. The mean residue levels of tetracycline were 503.67 μg/kg ±110.30 μg/kg, ciprofloxacin 22.4 μg/kg ± 5.20 μg/kg and 474.4 μg/kg ± 119.74 μg/kg for oxytetracycline respectively. The result shows multi-residues of antibiotics.

The sample 4 above tested shows positive for the two antibiotics tetracycline and ciprofloxacin residues. A total of 3 tissues (liver = 1; kidney = 1; muscle = 1) from 1 slaughtered cattle were screened for the presence of antibiotics residue. The result shows multi-residues of antibiotics. The mean residue levels of tetracycline were 49.7 μg/kg ± 6.14 μg/kg and 53.03 μg/kg ± 8.25 μg/kg for ciprofloxacin respectively.

It can be observed from the above samples 5 of 1 slaughtered cattle. A total of 3 tissues (liver = 1; kidney = 1; muscle = 1) were screened for the presence of drugs residue. Oxytetracycline were detected. The mean residue levels of Oxytetracycline were 227.2 μg/kg ± 16.45 μg/kg.

The result of analysis in sample 6 above shows positive for the two antibiotic tetracycline and ciprofloxacin residues respectively. A total of 3 tissues (liver = 1; kidney = 1; muscle = 1) from 1 slaughtered cattle were screened for the presence of antibiotics residue. The mean residue levels of tetracycline were 594 μg/kg ± 47.71 μg/kg and 49.6 μg/kg ± 6.42 μg/kg for ciprofloxacin respectively. The result shows multi-residues of antibiotics.

## 4. Discussion

Antibiotic use contributes to the emergence of drug resistant organisms, these important drugs must be used judiciously in both animal and human medicine to slow the development of resistance. The results of this study show the presence of residues of tetracycline, ciprofloxacin and oxytetracycline, which are some of the leading antimicrobials used in Nigeria, in all the tissues screened, with the highest concentrations found in the liver, followed by the kidney and then the muscle, with the lowest levels. This is incomparable with a study completed by Adesokan *et al*., (2013) which found that the excessive concentrations of antibiotics are within the muscular tissues, accompanied by the liver and kidney with lowest concentration. This finding further gives proof that abuse of antimicrobial use, such as when used on food animals meant for almost immediately slaughter, ends in residues in tissues obtained from table 3 above, out of 18 samples that were analyzed for tetracycline residues 15(83.3%) had detectable tetracycline residues, of those 15 samples, 5(83%) were from liver, 5(83%) were from kidney, and 5(83%) were from muscle. This study does not concur with findings of Muriuki *et al*., (2001) where out of the 250 samples that were analyzed for tetracycline residues, 114(45.6%) had detectable tetracycline residues. Of those 114 samples, 60(24%) were from liver, 35(14%) from kidney, and 19(7.6%) from muscle. Although the concentration levels obtained in this study were above the maximum residual limits (100) recommended (WHO/FAO, 1999) for some of the samples.

**Table 3:** Result of meat/organ with tetracycline, ciprofloxacin and oxytetracycline residue from Kano abattoir

| Sample Type | N | Tetracycline | Ciprofloxacin | Oxytetracycline | >MRLs |
| --- | --- | --- | --- | --- | --- |
| Liver<br>(22%) | 6 | 5 (83%) | 4 (67%) | 2 (33%) | 4 |
| Kidney | 6 | 5 (83%) | 4 (67%) | 2 (33%) | 0 (0%) |
| Muscles<br>(28%) | 6 | 5 (83%) | 4 (67%) | 2 (33%) | 5 |
| Total<br>(50%) | 18 | 15 (249%) | 12 (201%) | 6 (99%) | 9 |

Where: N is the number of samples, MRL is maximum residue limit

The result reported in figure 1 above comprising of liver = 1; kidney = 1; and muscle = 1 were screened for the presence of antibiotics residue. Tetracycline and ciprofloxacin were detected. The mean residue levels of tetracycline were 17.57 μg/kg ± 6.20 μg/kg and 82.77 μg/kg ± 12.60 μg/kg for ciprofloxacin respectively. The result shows multi-residues of antibiotics. This is consistent with finding of Muriuki *et al*., (2001) in Kenya with the mean tetracycline levels of cattle samples from the five slaughterhouses in the study were as follows: Athi River, 1,046 μg/kg; Dandora, 594 μg/kg; Ngong, 701 μg/kg; Kiserian, 524 μg/kg; and Dagoretti, 640 μg/kg. The mean levels of the detected tetracyclines were higher than the recommended maximum levels in edible tissues except in kidney of this present finding.

**Figure 1:**
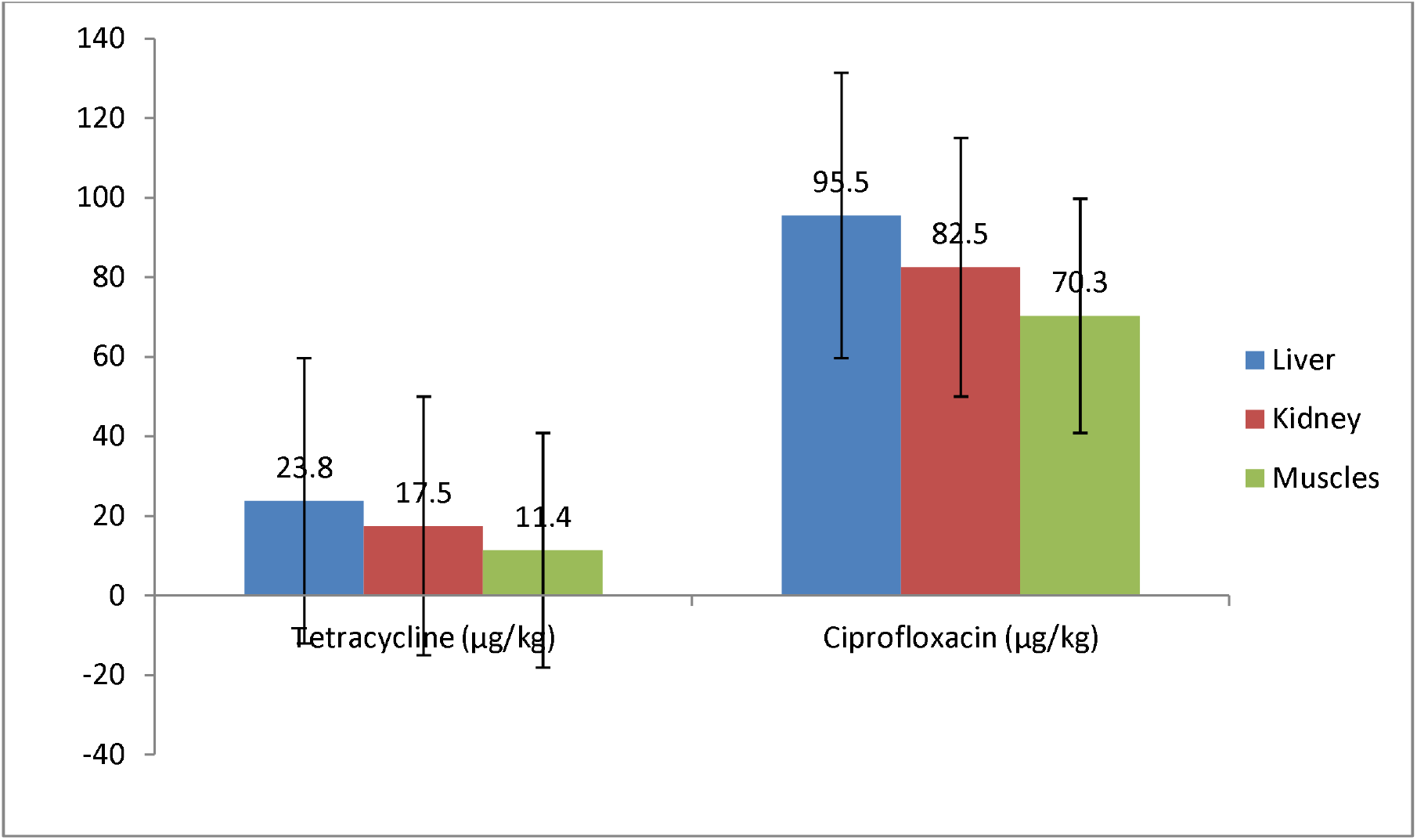
Mean distribution of Tetracycline and Ciprofloxacin residues in tissues of slaughtered cattle in sample 1.

It can be observed that from the figure 2 above comprising of liver = 1; kidney = 1; and muscle = 1 were screened for the presence of drugs residue. Only tetracycline was detected. The mean residue levels of tetracycline were high (575.37 μg/kg ± 73.23 μg/kg). The residue level in muscle is above permissible levels recommended by WHO/FAO, (1999). This is in contrast with work done by Kurwijila *et al*., (2006) on title: Investigation of the risk of exposure to antimicrobial residues present in marketed milk in Tanzania with mean residue of 404.72μg/kg ± 382.01 μg/kg.

**Figure 2:**
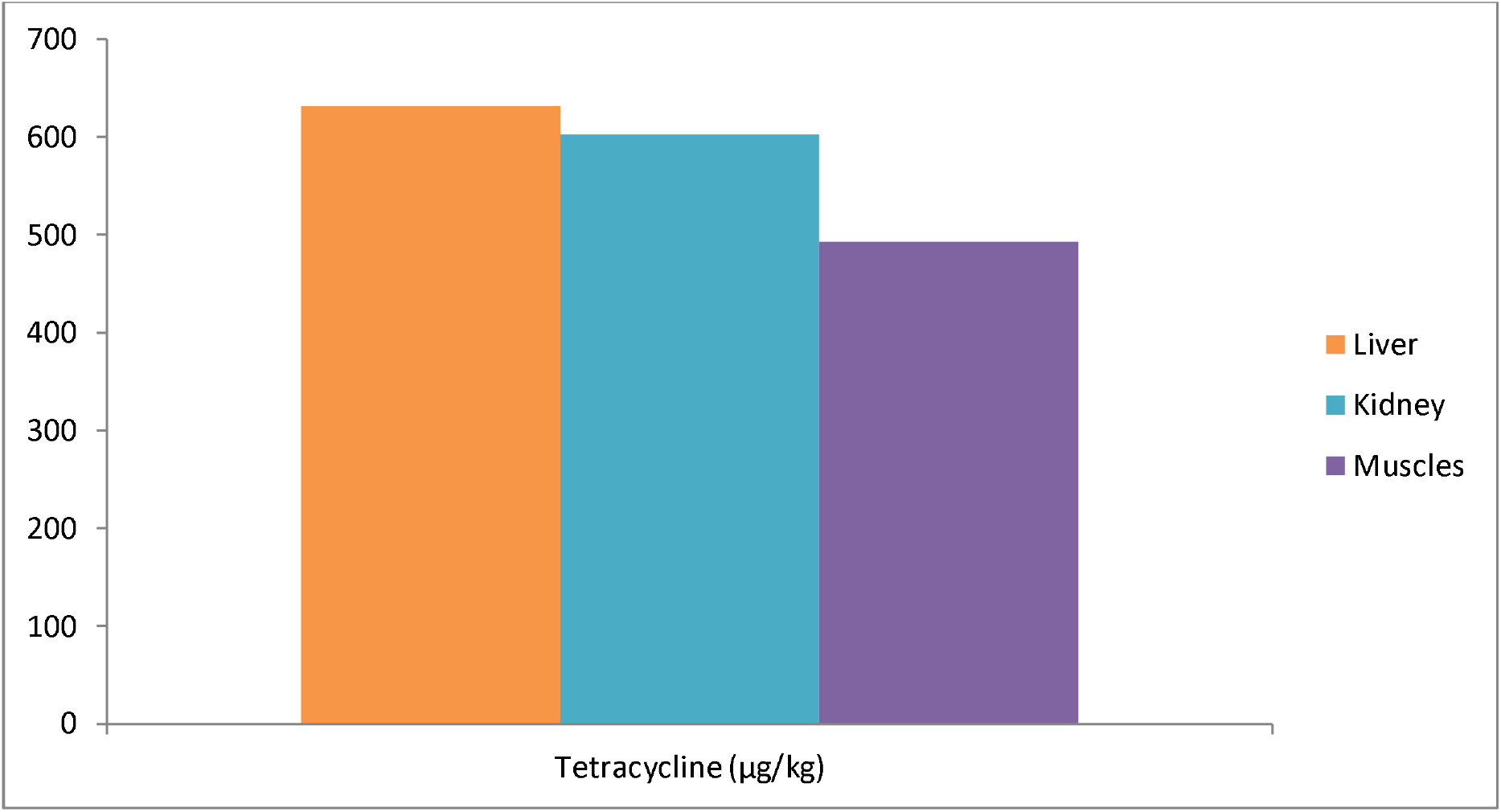
Mean distribution of Tetracycline residues in tissues of slaughtered cattle in sample 2.

Mean whilst, From the figure 3 above comprising of liver = 1; kidney = 1; and muscle = 1 were screened for the presence of antibiotics residue. Tetracycline, ciprofloxacin and oxytetracycline were detected. The mean residue levels of tetracycline were 503.67 μg/kg ±110.30 μg/kg, ciprofloxacin 22.4 μg/kg ± 5.20 μg/kg and 474.4 μg/kg ± 119.74 μg/kg for oxytetracycline respectively. The result shows multi-residues of antibiotics. This study disagrees with research by Sarker *et al*., (2018) where, out of the 160 samples that were analyzed for antibiotics ciprofloxacin residues. The percentage of the samples were 52% and 42% for liver and muscle respectively. The concentration levels obtained in this study were below the maximum residual limits recommended (WHO/FAO, 1999) except for the high concentration of Tetracyclin and oxytetracyclin in the muscles.

**Figure 3:**
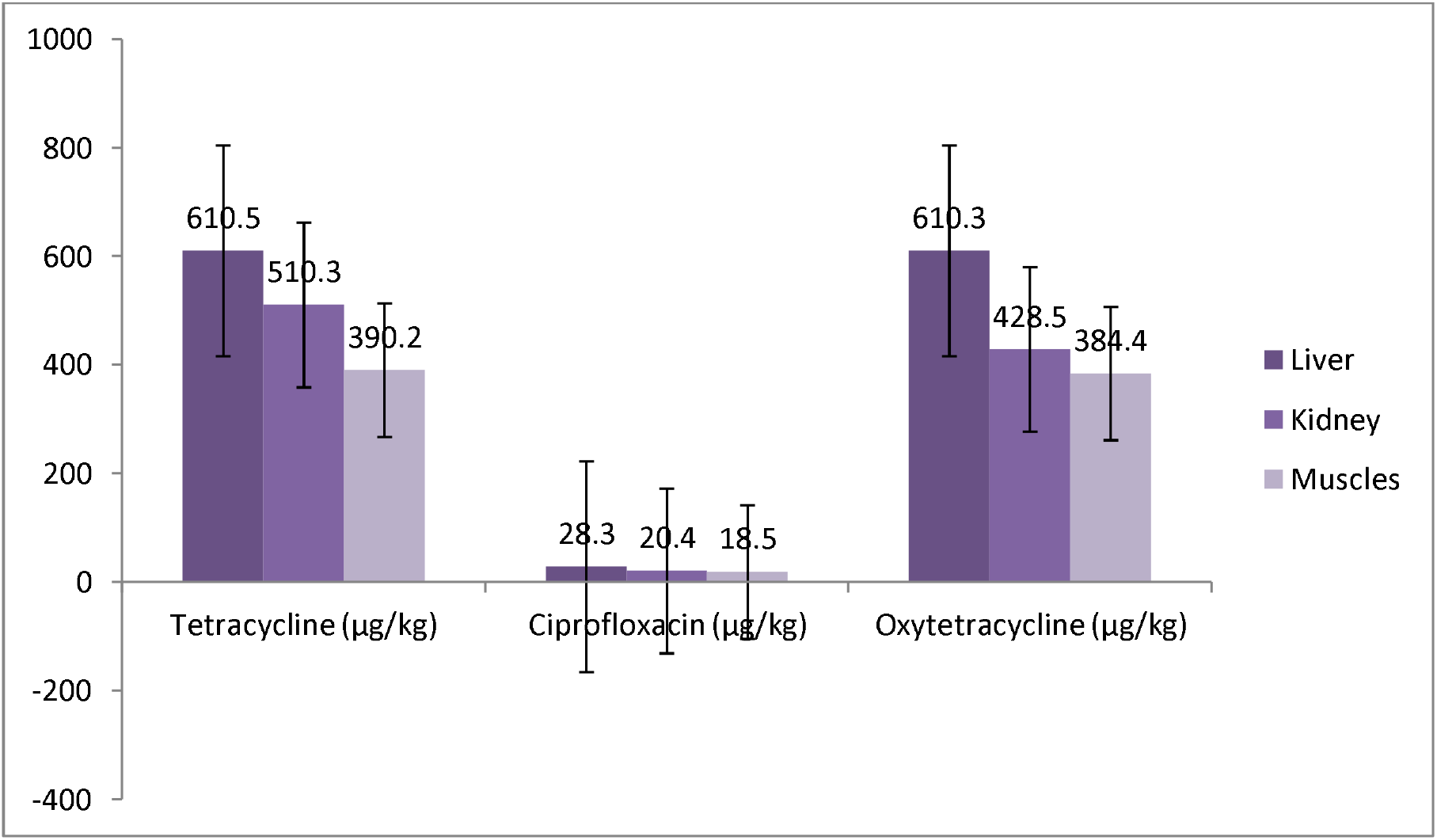
Mean distribution of Tetracycline, Ciprofloxacin and Oxytetracycline residues in tissues of slaughtered cattle in sample 3.

The figure 4 above detected two antibiotics tetracycline and ciprofloxacin residues. The mean residue levels of tetracycline were 49.7 μg/kg ± 6.14 μg/kg and 53.03 μg/kg ± 8.25 μg/kg for ciprofloxacin respectively. This concurs with a report by Olatoye and Ehinmowo, (2010), where out of the 180 samples that were analyzed for oxytetracycline residues, 98(54.44%) comprising of 48(80%) liver, 33(55%) kidney and 17(28.3%) muscle samples had detectable level of oxytetracycline residues. Out of the positive sample, 62(63.2%) had oxytetracycline residue at violative levels while 36(36.8%) had residues level below the WHO/FOA recommended MRLs for oxytetracycline in meat, liver and kidney with mean residue levels of 1197.0 μg/kg; 372.7 μg/kg and 51.80 in μg/kg in liver, kidney and muscles and 503.94 μg/kg, 492.07 μg/kg, 259.52 μg/kg for ciprofloxacin respectively.

**Figure 4:**
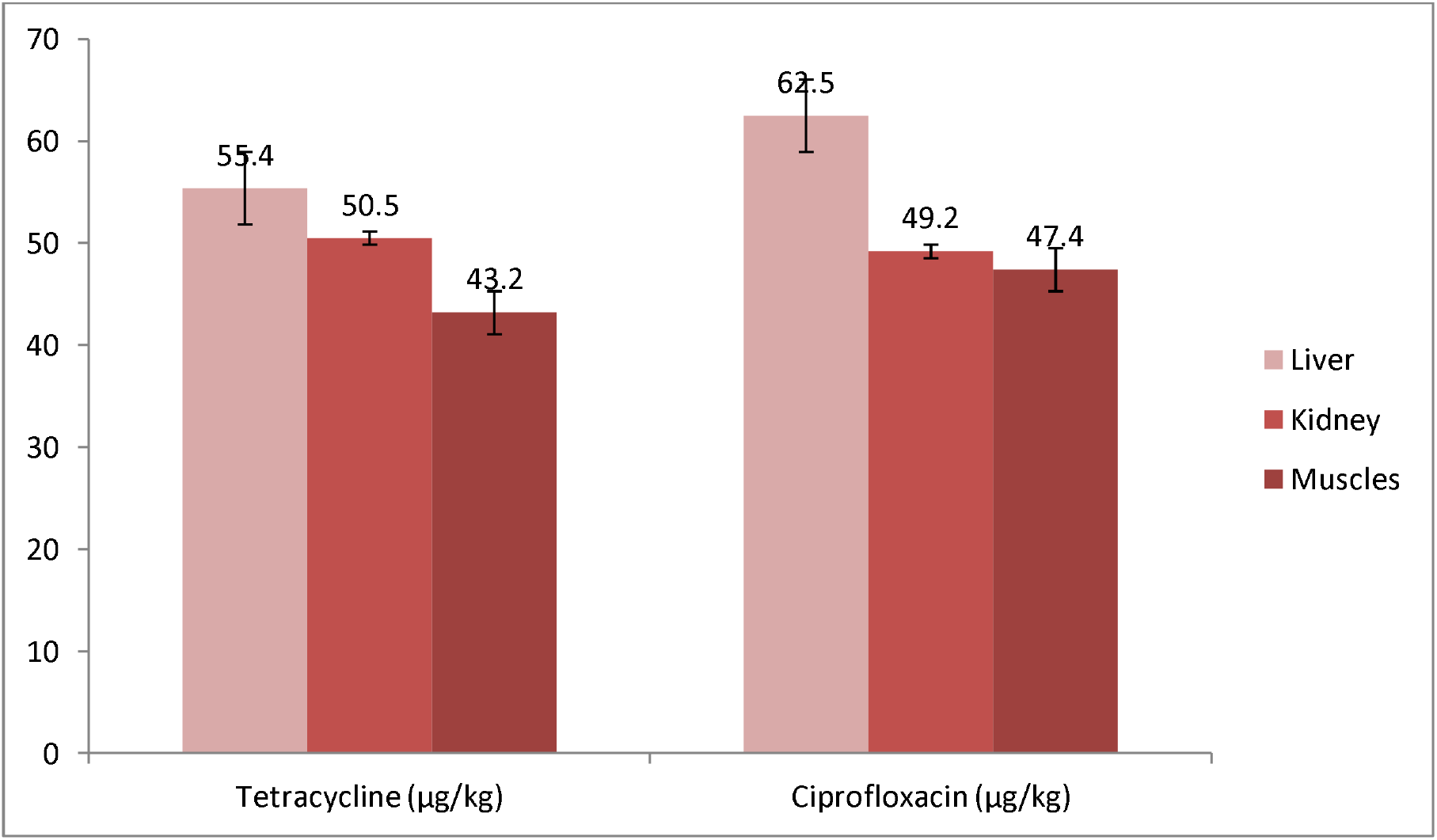
Mean distribution of Tetracycline and Ciprofloxacin residues in tissues of slaughtered cattle in sample 4.

It can be observed from figure 5 above comprising of liver = 1; kidney = 1; and muscle = 1 were screened for the presence of drugs residue. Oxytetracycline were detected. The mean residue levels were 227.2 μg/kg ± 16.45 μg/kg which is below the maximum permissible levels except in muscle. This is in agreement a study by Adesokan *et al*., (2013), in south-western Nigeria. The findings revealed residues of oxytetracycline (kidney: 9.47 μ/kg ± 3.24 μ/kg; liver: 12.73 μ/kg ± 4.39 μ/kg; muscle: 16.17 μ/kg ± 5.52 μ/kg). Although finding from these studies revealed that concentration of Oxytetracyclin in muscles where a little above the permissible limits.

**Figure 5:**
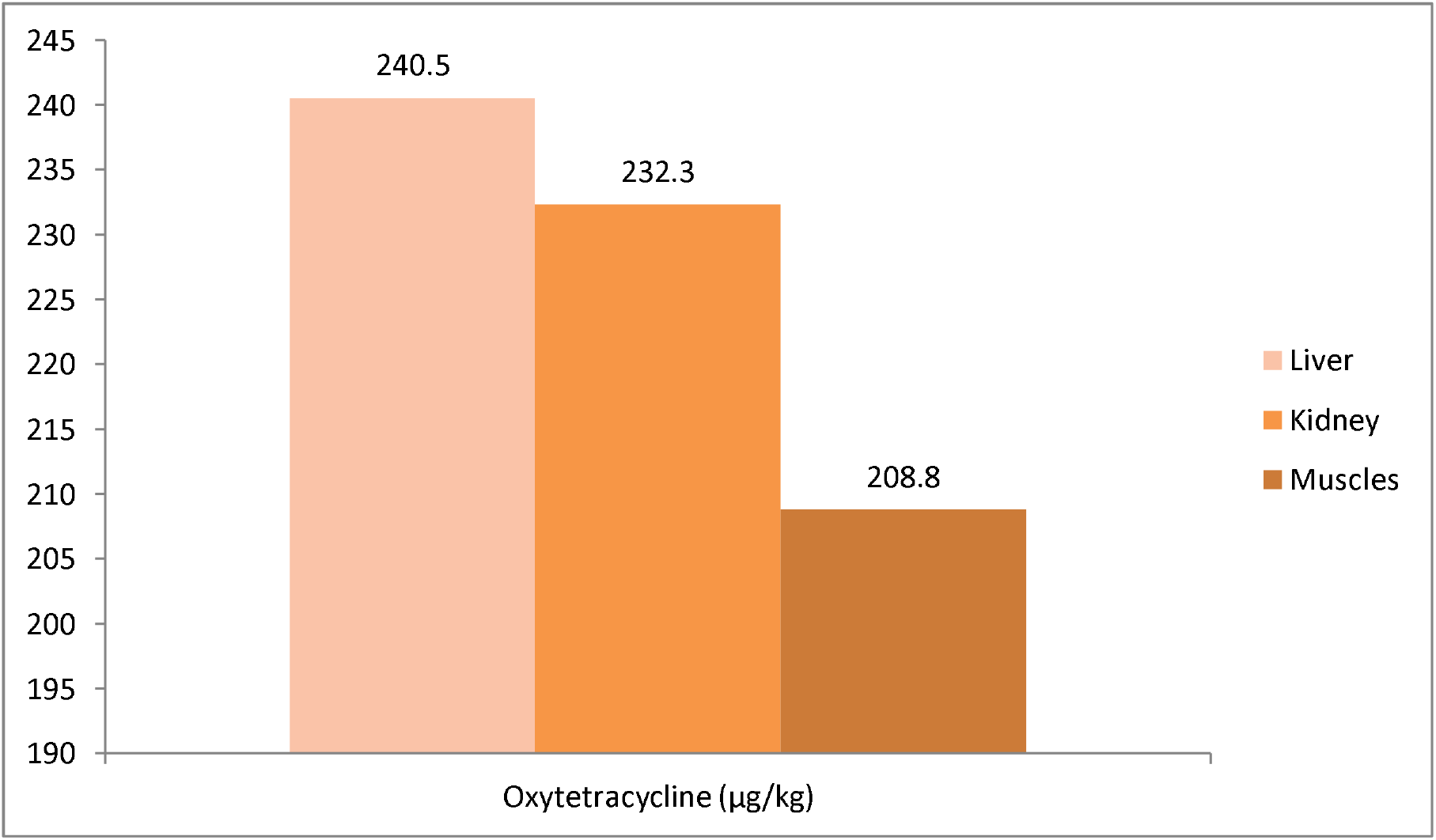
Mean distribution of oxytetracycline residues in tissues of slaughtered cattle in sample 5.

Results from figure 6 above detected two antibiotic tetracycline and ciprofloxacin residues respectively. This include liver = 1; kidney = 1; and muscle = 1 were screened for the presence of antibiotics residue. The result shows multi-residues of antibiotics. The mean residue levels of tetracycline were 594 μg/kg ± 47.71 μg/kg and 49.6 μg/kg ± 6.42 μg/kg for ciprofloxacin respectively. This is consistent with finding done by Ogara *et al*., (2001) in Nairobi slaughter house in Kenya except in kidney which is below permissible levels with mean residue of 366.8 μg/kg ± 421.7 μg/kg for tetracycline. Although, concentration of Tetracycline in this study was above the WHO recommended Maximum Residual Limits. The concentration level of Tetracycline in muscle samples in sample 2, oxytetracycline and tetracycline in muscles in sample 3, and tetracycline in muscles in samples 6 obtained in this study were above the maximum residual limits recommended (WHO/FAO, 1999). This is so within the light of bioaccumulation whilst these residues present in the animal tissues collect continuously over the lifespan of the individuals through extended intake. This is particularly of potential problem in Nigeria in which cattle are the most regularly ingested animals (Adetunji and Rauf, 2012; Gambo *et al*., 2010) and consumption of uncooked unpasteurized milk is normal among cattle owners (Cadmus *et al*., 2006), which can have a potentially bad impact on children, who are expected to consume more milk. Moreover, tetracycline, ciprofloxacin and oxytetracycline can input the human frame from other means which includes poultry meat, given the uncontrolled abuse of antibiotics amongst food animals in Nigeria.

**Figure 6:**
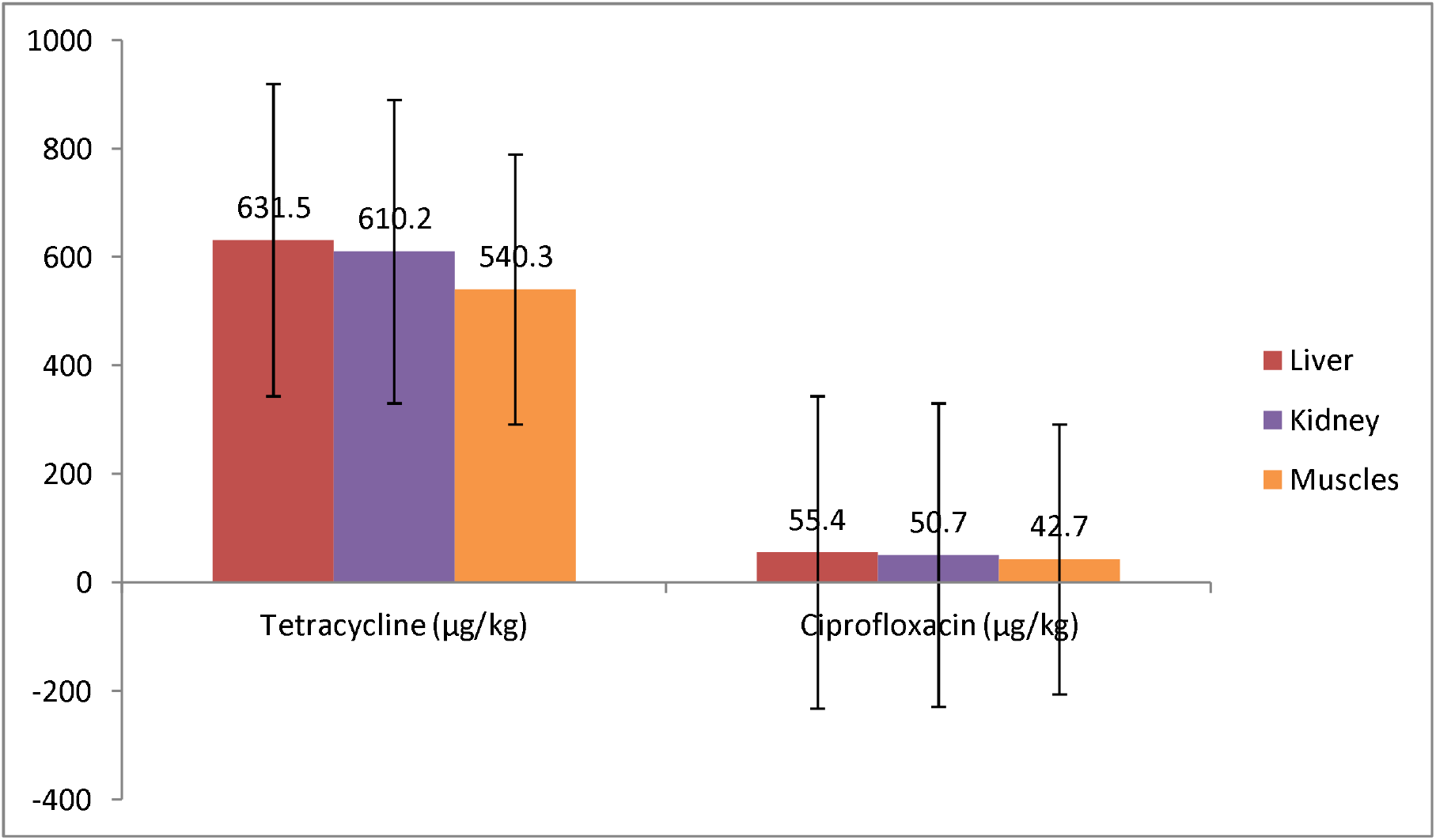
Mean distribution of Tetracycline and Ciprofloxacin residues in tissues of slaughtered cattle in sample 6.

The low levels detected in some of the tissues sampled could be attributed to the probably low doses of antimicrobials commonly administered by livestock traders in order to maximize the number of doses available, an assertion supported by an earlier report that nomadic herdsmen administer chemotherapeutic agents without veterinary prescription and most likely at incorrect dosages (Olatoye and Ehinmowo, 2010). In another study conducted in south-western Nigeria (Aibinu *et al*., 2007), strains of *E. coli* resistant to tetracycline and penicillin-G from both animal and human samples were reported. In general, reports from different parts of Nigeria have observed temporal trends in the prevalence of resistance amongst enteric organisms such as *E. coli* and *Shigella* (Okeke *et al*., 2005). This practice by livestock traders and herders therefore becomes an issue that needs urgent attention if efforts towards stamping out the menace of antimicrobial resistance in animals and humans.

## 5. Conclusion

The combination of several substances with diverse use as well as emission patterns, affecting a host of diverse endpoints in a plethora of exposed species in the vastly diverse ecosystems of the world, plus human health consideration, makes the derivation of a single quantitative antibiotic’s residue level in raw meat a daunting, if not impossible task. However, antibiotic traces have been found in the meat of cattle slaughtered for human consumption in Kano state. The findings in this study elucidated the importance of antibiotics usage in cattle and poultry and provided the quantitative analysis of the prevalence and levels of antibiotic residues. This study also provided quantitative data on the prevalence of oxytetracycline, tetracycline and ciprofloxacin residue in cattle meat that were being consumed in city of Kano, Nigeria. The results of this study revealed that greater proportion of the meat being consumed in Kano contained oxytetracycline and tetracycline residues above international food safety standards. The high proportion of cattle meat samples containing residues of antibiotics could be due to the misuse and lack of strict regulation and control of antimicrobial use Nigeria livestock production. This therefore requires urgent public health attention by the consumers and appropriate regulatory agencies in the country.

## 6. Recommendations

The misuse of antibiotics may result in the different health hazards. Thus, the reduction of antibiotic use constitutes a challenge for the world. In order to achieve such a reduction, the following nine steps should possibly be considered with regard to all antibiotics:

- Routine inspection should be adopted by environmental health officers to ensure drugs residues in animal product.
- Health education and sensitization of herdsmen and livestock industry should be established by environmental health officer for the negative impact of antibiotics residue on human health.
- The effective prevention of infectious diseases and the adoption of strict hygiene standards and rearing skills may reduce our need for antibiotics, particularly in the veterinary field.
- The use of alternatives to antibiotics, such as plant-derived antimicrobial substances and probiotics, may represent a promising option; vaccination against some bacterial diseases may be of great value in the near future.
- Strict national legislation must be passed in Nigeria to avoid the unnecessary use of antibiotics.
- National monitoring of antibiotic residues in foods and updating of the maximum permissible limits of these residues should be undertaken. In Table 1, we state the maximum limits on commonly observed antibiotic residues in foodstuffs.
- Avoid using antibiotics in the veterinary field without a veterinarian’s prescription o The heat treatment of meat, milk, and eggs may inactivate antibiotic contaminants in feedstuffs.
- The freezing of animal-derived foods may also contribute to the reduction of some antibiotic contamination.

